# Size-dependent segregation controls macrophage phagocytosis of antibody-opsonized targets

**DOI:** 10.1101/250373

**Authors:** Matthew H. Bakalar, Aaron M. Joffe, Eva M. Schmid, Sungmin Son, Marija Podolski, Daniel A. Fletcher

## Abstract

Macrophages protect the body from damage and disease by targeting antibody-opsonized cells for phagocytosis. Though antibodies can be raised against antigens with diverse structures, shapes, and sizes, it is unclear why some are more effective at triggering antibody-dependent phagocytosis than others. Here we quantitatively define an antigen height threshold that regulates phagocytosis of both engineered and cancer-specific antigens by macrophages. Using a reconstituted model of antibody-opsonized target cells, we find that phagocytosis is dramatically impaired for antigens that position antibodies >10 nm from the target surface. Increasing antigen height allows for co-localization of Fc receptors and the inhibitory phosphatase CD45 at the cell-cell interface, which reduces Fc receptor phosphorylation, and inhibits phagocytosis. Our work shows that close contact between macrophage and target cell is a requirement for efficient phagocytosis, suggesting that therapeutic antibodies should target short antigens in order to trigger Fc receptor activation through size-dependent physical segregation.

## INTRODUCTION

Controlled activation of immune cells is essential to protect the body from pathogens and diseased cells while limiting damage to healthy cells. Antibodies provide one way to control immune response by specifically targeting cells displaying foreign antigens for destruction by phagocytes and other innate immune cells. Macrophages use antibody-dependent cellular phagocytosis (ADCP) to destroy bacterial and fungal pathogens as well as virally infected self-cells. Recently, ADCP has also been shown to contribute significantly to anti-tumor immunity during monoclonal antibody (mAb) therapy (DiLillo et al., 2014; Erwig and Gow, 2016; Weiskopf and Weissman, 2015).

Over the past decade, the clinical use of mAbs to treat solid and hematological cancers has rapidly expanded. Humanized mAbs, including the anti-CD20 antibodies rituximab (Rituxan) and the anti-HER2 mAb trastuzumab (Herceptin), retarget Fc-receptor bearing immune cells against tumor cell targets, leading to cell death and clearance of malignant cells through myeloid cell mediated ADCP (Weiskopf and Weissman, 2015), as well as natural killer cell mediated antibody-dependent cellular cytotoxicity (ADCC) (Clynes et al., 2000). Recent studies have also suggested that the therapeutic mechanism of the checkpoint inhibitor mAb anti-CTLA-4 is dependent on Fc-dependent depletion of regulatory T cells by macrophages within the tumor (Simpson et al., 2013). While antibodies can be generated against almost any molecular target, the number of clinically successful antigens is surprisingly limited, consisting primarily of small cell-surface proteins (Shim, 2011). Further, different mAbs with the same target antigen can have widely varying modes of therapeutic action (Maloney et al., 2002).

What makes an antibody effective at stimulating ADCP? Biochemical properties of the antibody and target antigen, including binding affinity, isotype, and glycosylation, are known to be important (Raju, 2008). However, the role of physical properties in ADCP, specifically those of the antigen, remains unclear. Cell surface antigens can have diverse physical properties, including different structure, shape, and size. For example, members of the CEACAM family of cell-surface receptors are composed of 1-7 Ig-like domains, and their expression levels vary during cancer progression (Beauchemin and Arabzadeh, 2013). Whether antigen height affects ADCP could have important implications for antigen and epitope selection and for design of therapeutic mAbs, and how antigen height affects ADCP could provide insight into the molecular mechanisms that govern macrophage activation.

Antibody-dependent cellular phagocytosis is triggered by binding between the Fc region of IgG and macrophage Fcγ receptors (FcγRs). While the diversity of FcγRs and their affinity for IgG subclasses has been extensively studied (Nimmerjahn and Ravetch, 2005), the mechanism by which Fc binding to an FcγR leads to activation and phagocytosis is still under investigation. Unlike toll-like receptors, which dimerize upon ligand binding (Akira and Takeda, 2004), FcγR bind to IgG with one-to-one binding stoichiometry, and there is no known conformational change in FcγR upon binding (Lu et al., 2011). Instead, it has been suggested that clustering of multiple FcγRs on the fluid plasma membrane is necessary for receptor activation (Goodridge et al., 2012). This clustering is associated with phosphorylation of the FcγR immunoreceptor tyrosine-based activation motif (ITAM), which is controlled by a balance of Src family kinase (SFK) and tyrosine-phosphatase activity and is required for the activation of phagocytosis (Fitzer-Attas et al., 2000; Zhu et al., 2008).

Signaling through phosphorylated ITAMs occurs in macrophages as well as other immune cells, including T-cells, where T-cell receptor (TCR) phosphorylation similarly depends on a balance of SFK and tyrosine-phosphatase activity (McNeill et al., 2007; Zikherman et al., 2010). The kinetic segregation model of TCR signaling proposes that the phosphatases CD45 and CD148 are physically excluded from sites of TCR-pMHC binding due to their large extracellular domains, resulting in a local decrease in phosphatase activity around the TCR that leads to phosphorylation and activation of the receptor (Choudhuri et al., 2005; James and Vale, 2012; Varma et al., 2006). It has been suggested that a similar size-dependent mechanism of phosphatase segregation at close contacts between a macrophage and a target cell may trigger FcγR activation (Goodridge et al., 2012), but direct evidence is lacking. Furthermore, alternate mechanisms for signaling through FcγR have been reported, including establishment of a diffusion barrier by integrin binding (Freeman et al., 2016) and concentration of SFKs and FcγR into lipid micro-domains (Beekman et al., 2008; Katsumata et al., 2001).

Here we show that antibody-dependent phagocytosis and FcγR signaling is critically dependent on the height of an antibody above the target cell surface, and we exploit this size dependence to gain a better understanding of the molecular mechanisms that govern macrophage FcγR activation. Using a minimal reconstitution of the target cell surface and engineered antigens of variable height to isolate the signaling interactions between a macrophage and antibody-opsonized target, we find that physical segregation of large inhibitory phosphatases is necessary for FcγR phosphorylation and subsequent phagocytosis. Segregation – and consequently phagocytosis – are disrupted by antigens that are >10 nm from the target-cell surface. We find that this size dependence holds true not only for engineered antigens but also for the tumor-expressed CEACAM antigens. Using genome editing to reduce the size of the endogenous CD45 ectodomain, we show that a shorter phosphatase can disrupt phagocytosis of target particles even for antibodies that are bound close to the membrane.

Our results demonstrate that exclusion of large phosphatases is necessary for receptor activation and phagocytosis, suggesting that therapeutic mAbs intended to trigger ADCP will be most effective when targeting antigens within 10 nm from the cell surface. This size-dependent segregation mechanism may allow macrophages surveilling tissue to physically distinguish ‘casual’ contact with cells and particles that are not phagocytic targets from ‘close’ contact with those that are.

## RESULTS

### Reconstitution of a cell-like target particle for FcγR-mediated phagocytosis

In order to isolate the mechanism of FcγR signaling in macrophages, we reconstituted a minimal model of an antibody-opsonized target cell-surface *in vitro*. This model system consisted of glass micro-beads coated with a fluid supported-lipid bilayer (SLB) and antibody (Figure 1A). To bind antibodies close to the bilayer, we incorporated lipids with a biotinylated head-group into the membrane and incubated with a monoclonal anti-biotin IgG1 antibody (anti-biotin IgG). The biotin-bound antibody is able to diffuse fluidly on the membrane surface, emulating diffusion of a mammalian cell-surface antigen. To investigate phagocytosis of these cell-surface like target particles, we added them to an imaging chamber seeded with RAW 264.7 macrophage-like cells. At the point of contact between an opsonized target particle and a macrophage, we observed a striking enrichment of labeled anti-biotin IgG (Figure 1B), consistent with binding between the surface-bound antibody and macrophage FcγR and subsequent enrichment of the antibody-FcγR complex at the contact site. We observed little to no internalization of non-opsonized target particles but saw robust phagocytosis of target particles that were IgG-opsonized, including internalization of multiple beads per cell (Figure 1C).

To quantify phagocytosis, we used confocal microscopy to image the internalization of target particles at single-cell resolution after incubation of beads and macrophages for 20 minutes. Total SLB fluorescence within a cell was used as a proxy for the amount of phagocytosis, and an automated image-segmentation routine was developed to quantify the per-cell fluorescence of the internalized lipid (Figure 1D and E, and Experimental Procedures ‘Microscopy assay of phagocytosis’). Opsonization with anti-biotin IgG is necessary and sufficient for phagocytosis, as evidenced by the lack of internalization in the absence of antibody or biotinylated lipid. Phagocytosis increased with increasing concentrations of anti-biotin IgG and could be blocked by incubating macrophages with antibodies (Fc block) against CD16 (FcγRIII) and CD32 (FcγRIIB), demonstrating that internalization of the opsonized SLB-coated beads is FcγR specific (Figure 1E).

### Production of size-variant antigens based on a synthetic FNIII domain

Cell-surface molecules often contain repeats of common domains such as Ig and FNIII (A F Williams and Barclay, 1988). To model antigens of different heights on our target particles, we developed a family of size-variant proteins based on repeats of a synthetic FNIII domain (Fibcon), which has a size of 3.5 nm (Figure 1F) (Jacobs et al., 2012). Each protein consists of repeats of the Fibcon domain with no inter-domain linker. We named the repeat proteins Fib1L, Fib3L, Fib5L, and Fib7L to denote the number of repeated domains. An N-terminal His-tag on each protein enables it to bind nickel-chelating lipids incorporated into the bead SLB, and proteins were expressed and purified in E. coli to exclude glycosylation.

In an fully extended configuration, the Fibcon-family proteins have maximum lengths of 3.5 nm, 10.5 nm, 17.5 nm, and 24.5 nm respectively. To quantify their average extension when bound to an SLB via the His-tag, we used a single-axis fluorescence localization method to measure the distance between the protein N-terminus and the SLB (Experimental Procedures). Our measurements show that the Fibcon repeat proteins bind to the bilayer in an upright configuration, with measured heights of 5.0 +/− 0.40 nm, 8.9 +/− 0.34 nm, 11.0 +− 0.8 nm, and 12.2 +/− 0.64 nm (Figure 1F). As expected for a semi-flexible polymer, the measured height diverges from the maximum length of the extended configuration as the number of Fibcon domains in the protein increases.

To turn the Fibcon-family proteins into antigens with identical well-defined epitopes, we incorporated an N-terminal YBBR tag for site-specific enzymatic modification and used SFP synthase to enzymatically couple biotin-CoA to the proteins (Figure S1A and B). The result was a family of size-variant antigens that diffuse freely on the minimal target particles and can be targeted by anti-biotin IgG (Figure 1G).

**Figure 1.**
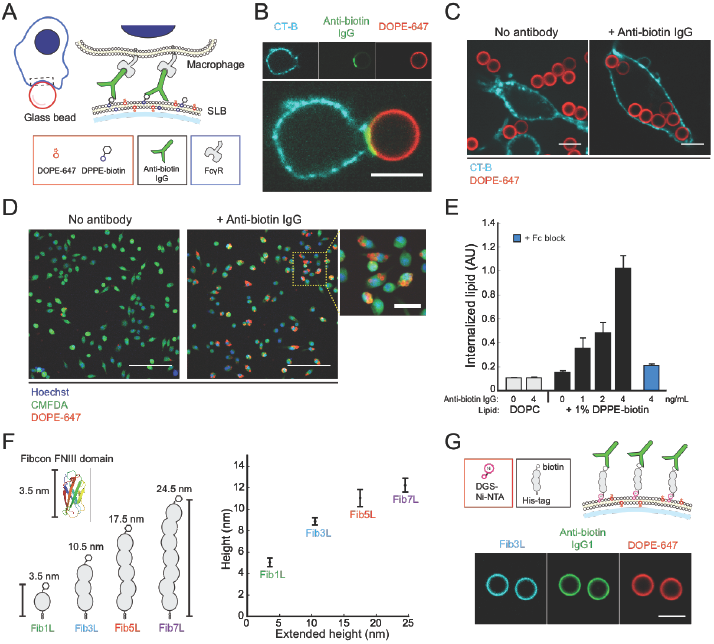
Reconstitution of a cell-like target particle for FcγR-mediated phagocytosis. (A) Target particles assembled *in vitro* from glass beads coated in a fluid supported lipid bilayer (lipid composition: DOPC, 0.2% DOPE-647, up to 2% DPPE-biotin). Anti-biotin IgG in solution binds fluidly to the lipid surface through interaction between the antigen-binding region of IgG and the biotin head-group of DPPE-biotin. Contact between a macrophage and a target particle leads to binding between FcγRs on the macrophage surface and the Fc region of anti-biotin IgG. (B) Confocal fluorescence images (60x) of a RAW 264.7 macrophage-like cell at a contact interface with a 6.46 μm target particle. The macrophage membrane (cyan) is labeled with cholera-toxin B 555, and the target particle membrane (red) contains fluorescent DOPE-647. Scale bar is 5 μm. (C) 3.78 μm target particles were added to an imaging well containing RAW 264.7 cells. Addition of target particles containing only lipid (left). Addition of target particles pre-incubated with 4 ng/mL anti-biotin IgG (right). (D) Representative confocal fluorescence images (20x) of a field-of-view (FOV) from the imaging-based phagocytosis assay. After gently washing, images reveal very few beads internalized for target particles formed without anti-biotin IgG (left). Target particles bound with anti-biotin IgG are internalized and multiple beads per cell are visible within the macrophages (right). Cells are labeled with 0.5 μM CellTracker Green (CMFDA) and 10 μg/mL Hoechst 33342. Scale bar is 100 μm for large field of view (FOV), 25 μm for zoom-in. (E) Quantification of internalized fluorescence from internalized beads. Error bars are standard error across 6 independent wells. For each well, internalized lipid is an average quantification of N > 250 cells. f, A size-variant antigen family is constructed from repeats of the Fibcon synthetic FNIII domain (pdb 3TEU). Proteins in the family are named Fib1L, Fib3L, Fib5L, and Fib7L, and when extended have heights of 3.5 nm, 10.5 nm, 17.5 nm, 24.5 nm respectively (left). The distance between the lipid bilayer and the N-terminus of the antigen was measured using a one-dimensional fluorescence localization method (right) (see Experimental Procedures). The N-terminal height above the bilayer for Fib1L, Fib3L, Fib5L, and Fib7L was quantified at 5.0 +/− 0.40 nm, 8.9 +/− 0.34 nm, 11.0 +− 0.8 nm, and 12.2 +/− 0.64 nm respectively. g, Fibcon proteins with a C-terminal His-tag were N-terminally labeled with biotin to construct synthetic protein antigens that bind fluidly to a SLB coated bead containing 0.8% DGS-Ni-NTA lipid (top). Confocal fluorescence images (20x) of Fib3L antigen bound and anti-biotin IgG opsonized target particles (bottom). Scale bar is 5 μm.

### Phagocytosis of antibody-opsonized target particles is antigen-height dependent

To determine the impact of antigen height on phagocytosis, we quantified phagocytosis of target particles (3.78 μm) bound with biotinylated Fib1L, Fib3L, Fib5L, and Fib7L protein antigens opsonized with anti-biotin IgG (Figure 2A and Figure S1D). Strikingly, we observed decreasing phagocytosis with increasing antigen height; macrophages efficiently internalized beads coated with Fib1L antigen, while phagocytosis was significantly impaired against Fib3L antigen and nearly absent for Fib5L and Fib7L antigen coated beads (Figure 2B). Using flow-cytometry, we confirmed that equal concentrations of antibody were binding to each target particle (Figure S1C), suggesting equal concentrations of antigen. Therefore, differences in Fibcon-family antigen lengths – not antibody surface concentration – were responsible for the observed decrease in phagocytosis.

Since phagocytosis is known to be modulated by antibody concentration (Ben M’Barek et al., 2015), we investigated whether the observed antigen height-dependent phagocytosis was unique to one concentration. Using flow cytometry, we assayed phagocytic efficiency across a range of anti-biotin IgG concentrations for each antigen. While all antigen heights show increased phagocytosis with increasing concentrations of anti-biotin IgG, the data reveals a consistent size-dependent suppression of phagocytosis, with dramatically reduced sensitivity as a function of antibody concentration for Fib5L and Fib7L (Figure 2C). Our data suggest that short antigens promote efficient phagocytosis, while antigens that are > 10 nm have severely diminished phagocytic capacity.

### Phagocytosis of CEACAM antibody opsonized target particles is antigen-height dependent

We next explored whether antigen height dependence is unique to our synthetic proteins or a more general property of antibody-dependent phagocytosis. To address this, we used members of the CEACAM family of cell-surface proteins, which are associated with tumor progression and include both short and tall antigens (Beauchemin and Arabzadeh, 2013). We expressed and purified full-length CEACAM5 (CEA-FL, 28.0 nm estimated height) and a truncated version of CEACAM5 consisting of only the N-terminal domain (CEA-N, 4.0 nm) (Figure 2D and Figure S2A) (Korotkova et al., 2008). We selected a pan-CEACAM IgG1 antibody (anti-CEA IgG) that binds directly to the N-terminal domain of CEACAM5 (Figure S2B) so that target-cell membranes coated with CEA-FL or CEA-N could be opsonized with the same antibody as our synthetic antigens.

Consistent with our Fibcon-family antigen experiments, we found that the short CEA-N was efficiently internalized, while phagocytosis of the long CEA-FL was significantly reduced (Figure 2E, F). Using flow-cytometry, we again confirmed that equal concentrations of antibody were bound to CEA-FL and CEA-N coated target particles (Figure S2C). We assayed phagocytic efficiency across a range of concentrations of anti-CEACAM and observed that while phagocytosis increases with increasing antibody concentration for both antigens, the increase is significantly greater for CEA-N (Figure 2G). We conclude that antibody-dependent phagocytosis of CEACAM-family antigens is height-dependent, similar to our Fibcon-family antigens. Confocal images of cells acquired during the phagocytosis reveal that multiple beads-per-cell were internalized for CEA-N target beads, while CEA-FL target beads remain bound to the cell surface but not internalized. (Figure S2D).

**Figure 2.**
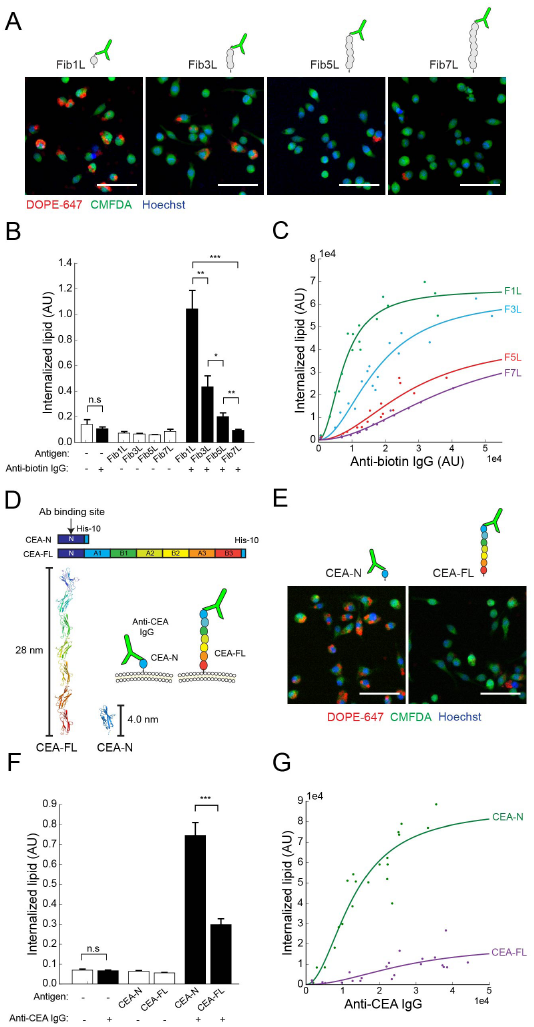
Phagocytosis of antibody-opsonized target particles is antigen-height dependent. (A) Representative confocal fluorescence images (20x) of phagocytosis of target particles bound with biotinylated Fib1L, Fib3L, Fib5L, and Fib7L protein antigen and opsonized with 4 ng/mL anti-biotin IgG. Cells are labeled with 0.5 μM CellTracker Green (CMFDA) and 10 μg/mL Hoechst 33342. Scale bar is 50 μm. (B) Microscopy quantification of phagocytosis for Fib1L, Fib3L, Fib5L, and Fib7L target particles. Error bars represent standard error across 9 independent wells. For each well, internalized lipid is an average quantification of N > 330 cells. P-values are paired Student’s T-test, where *P < 0.05, **P < 0.01, ***P < 0.001. (C) Flow-cytometry quantification of phagocytosis for Fib1L, Fib3L, Fib5L, and Fib7L targets for increasing anti-biotin IgG concentrations. Each data point corresponds to an independent well with N > 8000 cells. For each data point, IgG fluorescence intensity (anti-biotin IgG, Alexa Fluor 488) was pre-measured via flow cytometry from a sample of target particles (N > 3500 beads). Each set of data points is fit with a hill equation with a coefficient of 2 using the equation f(x) = (ymax*x^2^) / (kd + x^2^). (D) Full-length CEACAM5 (CEA-FL, 28.0 nm) and a truncated version of CEACAM5 consisting of the N-terminal domain (CEA-N, 4.0 nm). A pan-CEACAM IgG1 antibody (anti-CEA IgG) (D14HD11) binds directly to the shared N-terminal domain. (E) Representative confocal fluorescence images (20x) of phagocytosis of target particles bound with CEA-N and CEA-FL antigen opsonized with 4 ng/mL anti-CEA IgG. Cells are labeled with 0.5 μM CellTracker Green (CMFDA) and 10 μg/mL Hoechst 33342. Scale bar is 50 μm. (F) Quantification of phagocytosis for CEA-N and CEA-FL target particles from the microscopy assay. Error bars are standard error across 9 independent wells. For each well, internalized lipid is an average quantification of N > 420 cells. P-values are paired Student’s T-test, where ***P < 0.001. (G) Flow-cytometry quantification of phagocytosis for CEA-N and CEA-FL targets across a range of bound anti-CEA IgG concentrations. IgG fluorescence intensity (anti-CEA IgG, PE) was pre-measured via flow cytometry from a sample of beads (N > 3500 beads). Each set of data points is fit with a hill equation with a coefficient of 2 using the equation f(x) = (ymax*x^2^) / (kd + x^2^).

### Visualization of early Fc receptor signaling prior to phagocytosis

Our experiments with variable height antigens show a consistent and significant decrease in phagocytosis with increasing antigen height, indicating either a reduction in activation of the FcγR or a reduction of signal transduction downstream of the receptor. The first signaling event after FcγR-antibody binding is phosphorylation of the receptor’s ITAM motif by Src family kinases (SFK), followed by recruitment of Syk kinase to the ITAM via its tandem SH2 domains (Crowley et al., 1997). To test whether the antigen height-dependent phagocytosis we observed is linked to changes in ITAM signaling, we imaged the distribution of phosphotyrosine on cells fixed during phagocytosis of a target particle with a short antigen, Fib1L. Visualization of nascent phagocytic cups by immunofluorescence revealed enrichment of ITAM tyrosine phosphorylation, while for Fib7L target particles the increase in phosphotyrosine signal was notably absent at contact sites (Figure S3A).

To image the dynamics of ITAM phosphorylation in greater detail, we developed a live-cell protein-based ITAM phosphorylation sensor that is specific to phosphorylated ITAM (pITAM). The pITAM sensor is formed from the tandem SH2 domains of Syk kinase, which are known to mediate binding of Syk to the pITAM (Turner et al., 2000), and an N-terminal fusion with mCherry (Figure 3A and S3B, Supplementary Video 2). We established a stable RAW 264.7 cell line expressing pITAM sensor under the control of a constitutive weak promoter UBC to prevent competition with endogenous Syk localization and to reduce signal background (Qin et al., 2010). We replaced the SLB-coated target particles, which geometrically limit our ability to spatially resolve FcγR organization, with an SLB-coated coverslip (emulating the target surface) that permits high-resolution imaging in Total Internal Reflection Fluorescence (TIRF) microscopy (Figure 3B). Upon contact between cells expressing the pITAM sensor and a target surface coated with anti-biotin IgG opsonized Fib1L, we found that the sensor is rapidly recruited from the cytoplasm to antibody-FcγR clusters at the membrane interface (Figure 3C top, and S3C). The pITAM sensor dissociates from these clusters within seconds upon addition of the Src kinase inhibitor PP2, consistent with sensor-specificity for phosphorylated ITAM (Figure S3D).

### Fc receptor phosphorylation decreases with increasing antigen height

We next asked whether phosphorylation of the FcγR ITAM changes systematically with antigen height. To answer this, we used TIRF microscopy to quantify membrane-localized sensor fluorescence intensity at the macrophage-contact interface after cell spreading on a planar SLB coated with antibody-opsonized Fibcon-family antigens. We found that the level of pITAM sensor recruitment decreases significantly between Fib1L and Fib3L antigen, with recruitment dropping to near background levels for Fib5L and Fib7L (Figure 3C, D). We also noted that while anti-biotin IgG clusters at both Fib1L and Fib7L opsonized surfaces, there was a slight decrease in anti-biotin IgG concentration within these clusters for Fib7L antigen (Figure 3E).

Clusters of antibody-FcγR can be seen in the TIRF microscopy movies of the macrophage-opsonized surface trafficking inward from the periphery of the cell (Supplementary video 1), consistent with interactions between FcγR and the actin cytoskeleton that have been previously described (Freeman et al., 2016; Jaumouillé and Grinstein, 2011). Interestingly, the decrease in pITAM sensor recruitment with increase in antigen height was still evident in Latrunculin A treated cells, suggesting that interactions between FcγR and the cortical actin cytoskeleton is not necessary for receptor phosphorylation (Figure 3F). We also observed a decrease in FcγR ITAM phosphorylation in cells interacting with tall CEA-FL relative to short CEA-N antigen (Figure 3G), indicating that size-dependent phosphorylation of FcγR ITAMs is not unique to a single family of antigens.

**Figure 3.**
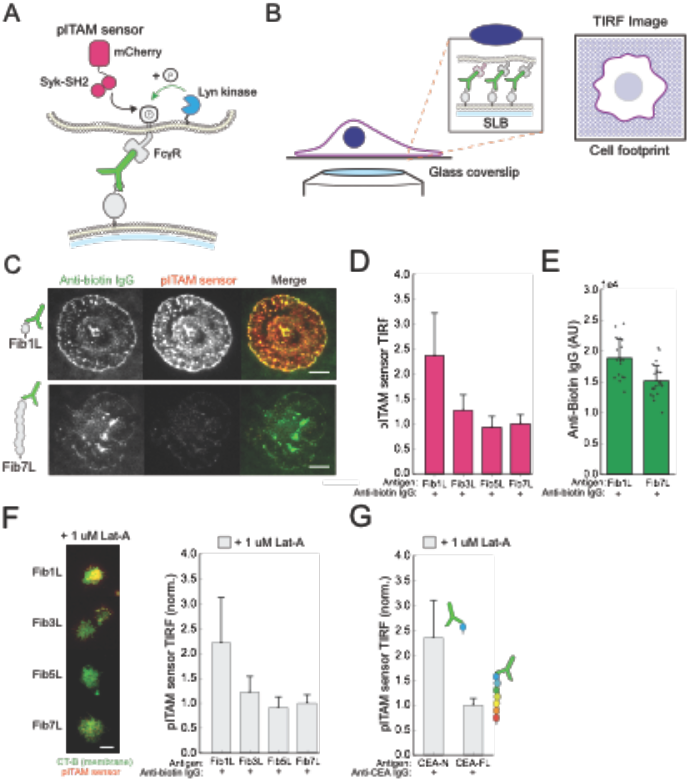
Fc receptor phosphorylation decreases with increasing antigen height. (A) A live-cell sensor of ITAM phosphorylation (pITAM sensor). The sensor consists of a N-terminal mCherry fluorescent protein flexibly linked to the tandem-SH2 domains of Syk kinase. Upon phosphorylation of FcγR ITAM by Src family kinases, the sensor protein is recruited to the phosphorylated ITAM through the tandem-SH2 domains. (B) TIRF microscopy of the interface between a macrophage and an antibody opsonized planar supported lipid bilayer enables high-resolution visualization of protein spatial organization at the contact site. (C) TIRF microscopy (100x) images of the contact interface between a macrophage and a supported lipid bilayer bound with Fib1L (top) or Fib7L antigen and opsonized with anti-biotin IgG. Scale bar is 10 μm. (D) Quantification of TIRF signal from pITAM sensor across the membrane-contact area at macrophage-SLB contact sites for Fib1L, Fib3L, Fib5L, and Fib7L bound SLBs. Error bars are standard deviation over N > 180 cells. (E) Quantification of mean TIRF signal from anti-biotin IgG (Alexa Fluor 488) from high-intensity clusters within the membrane-contact area at macrophage-SLB contact sites for Fib1L and Fib7L bound SLBs. Error bars are standard deviation from N > 18 cells. (F) TIRF images of anti-biotin IgG (green, Alexa Fluor 488) and pITAM sensor (red) at macrophage-SLB contacts where cells are treated with 1 μM Lat-A to depolymerize the actin cytoskeleton (left). Quantification of TIRF signal from pITAM sensor across the membrane-contact area at macrophage-SLB contacts for cells treated with Lat-A shows pITAM recruitment for Fib1L, and decreased recruitment for taller antigens, in the absence of a macrophage actin-cytoskeleton (right). Scale bar is 10 μm. (G) For SLBs bound with CEA-N and CEA-FL antigen and opsonized with anti-CEA IgG, quantification of TIRF signal from pITAM sensor across the membrane-contact area in the presence of Lat-A shows increased pITAM recruitment for CEA-N vs CEA-FL antigen. Error bars are standard error over 3 independent wells with mean intensity computed from N > 200 cells for each well.

### Antigen height alters antibody-FcγR affinity

A decrease in phosphorylation with height could reflect changes in local receptor clustering. Since total concentration of antibody on a target particle can modulate phagocytosis (Figure 1E, 2C, 2G), we asked whether there was a difference in effective concentration of FcγR within clusters for antigens of different heights. To test if the concentration of antibody-FcγR complex differed between Fib1L and Fib7L antigens, we measured the average intensity of anti-biotin IgG within these clusters (Figure 3E). We found that for a Fib7L antigen, the average concentration of clustered anti-biotin IgG was lower compared to the Fib1L antigen. This result could be explained by a size-dependent decrease in receptor-ligand affinity due to changes in binding entropy that is predicted physically for cadherins and has been observed at the T-cell immunological synapse (Milstein et al., 2008; Wu et al., 2011).

To determine if reduced receptor density was due to a decrease in receptor-ligand affinity with antigen height in the absence of other factors involved in cluster formation, we formed giant plasma membrane vesicles (GPMVs) from detached cellular-membrane blebs. These GPMVs have a lipid and membrane protein composition similar to the plasma membrane, but lack a cortical actin cytoskeleton and membrane cortex attachments. We isolated GPMVs from RAW 264.7 cells and added them to SLBs with antibody-opsonized Fibcon-family antigens (Figure 4A). The GPMVs settled onto the SLBs and formed planar footprints, with anti-biotin IgG bound to FcγR clearly enriched at the interface (Figure 4B). We quantified the ‘enrichment index’ of the anti-biotin IgG by taking the intensity ratio between regions beneath the GPMVs and regions of the SLB background. Since bound complexes become trapped at the interface, the ‘enrichment index’ provides a metric of relative receptor-ligand affinity (Schmid et al., 2016). Although we observed significant enrichment of anti-biotin IgG at the interface for each antigen, consistent with FcγR binding, the data revealed a consistent decrease in receptor-ligand affinity with increasing antigen height (enrichment index: Fib1L = 9.60 +/− 2.30, Fib3L = 8.31 +/− 1.44, Fib5L = 7.26 +/− 1.33, Fib7L = 6.82 +/− 9.99) (Figure 4C).

Since both receptor concentration and ITAM phosphorylation are decreased with antigen height, could lower receptor-ligand affinity alone account for our observation of reduced activation? Using images of macrophage ITAM phosphorylation on antibody-opsonized Fib1L and Fib7L surfaces, we plotted anti-biotin IgG intensity against pITAM sensor recruitment on a single-cell basis (Figure 4D). Despite cell-to-cell variations, the data shows that pITAM sensor recruitment to Fib1L is significantly higher than to Fib7L across the entire range of anti-biotin IgG concentration found in the wild-type macrophage population, suggesting that changes in receptor concentration alone cannot explain the decreased phosphorylation for tall antigens. As a further check, we renormalized the antibody concentration axis of our plot of Fibcon-family antigen-coated particle phagocytosis (Figure 2C) by the enrichment index for each antigen (Figure S1E). Interestingly, the renormalization caused the Fib5L and Fib7L trends to collapse onto a similar curve, suggesting that the difference in phagocytic efficiency between these antigens may be explained by affinity alone. However, the phagocytic response of the Fib1L and Fib3L remain significantly different, with the strongest response observed for the shortest Fib1L antigen, indicating a clear dependence of phagocytosis on antigen size rather than affinity.

**Figure 4.**
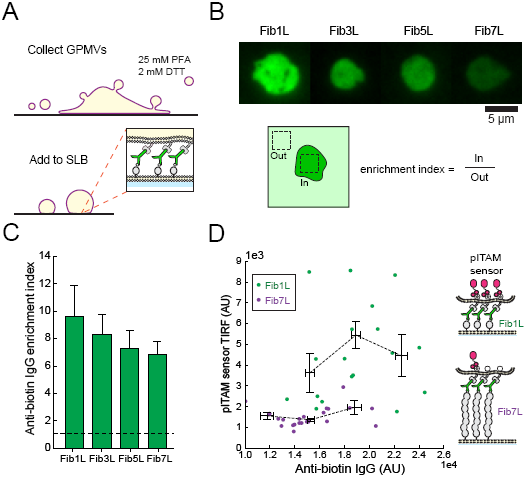
Antigen height alters antibody-FcγR affinity. (A) Giant plasma membrane vesicles (GPMVs) are formed by treating adhered macrophages with 25 mM PFA and 2 mM DTT to induce membrane blebbing and vesiculation, and collected from solution to recover vesicles with lipid and protein composition derived directly from RAW 264.7 cells. GPMVs dropped onto an opsonized supported lipid bilayer triggers binding between FcγR in the GPMVs and antibody on the SLB, adhering the GPMVs to the bilayer. (B) TIRF microscopy (100x) images of GPMV-SLB contacts at anti-biotin IgG opsonized SLBs for Fib1L, Fib3L, Fib5L, and Fib7L show a decrease in anti-biotin IgG (green, Alexa Fluor 488) intensity at the contact site with increasing antigen height. Anti-biotin IgG enrichment is calculated as the ratio of intensity within the GPMV-SLB contact site (in) and outside the contact site (out). Scale bar is 5 μm. (C) Quantification of anti-biotin IgG enrichment at GPMV-SLB contacts for Fib1L, Fib3L, Fib5L, and Fib7L bound SLBs. (D) Quantification of pITAM sensor intensity in TIRF as a function of anti-biotin IgG at single GPMV contacts for Fib1L and Fib7L bound SLBs. Across similar densities of anti-biotin IgG (from 1.5 – 1.9 AU), pITAM sensor TIRF is greater for Fib1L than for Fib7L antigen, showing enhanced phosphorylation at similar FcγR local densities.

### Phosphatase exclusion from antibody-FcγR clusters is antigen-height dependent

Phosphorylation of the FcγR ITAM is reversed by receptor-tyrosine phosphatases CD45 and CD148 (Zhu et al., 2008). To investigate how inhibitory phosphatases are spatially organized on the membrane relative to enriched antibody-FcγR, we used labeled antibodies to image the distribution of CD45 at the interface between a macrophage and an SLB bound with antibody-opsonized Fibcon-family antigens. TIRF images revealed striking segregation of anti-biotin IgG from CD45 when macrophages engaged with a Fib1L opsonized surface (Figure 5A). In contrast, we observed no segregation of CD45 from anti-biotin IgG clusters on a Fib7L opsonized surface.

We next asked whether CD45 was physically excluded from close membrane-membrane contacts formed by antibody-FcγR binding due solely to steric interaction between its large extracellular domain and the cell membranes. To quantify the ability of the membrane-interface to exclude large proteins, we again turned to GMPVs and generated interfaces between GPMVs and planar SLBs for each of the Fibcon-family antigens. TIRF images of the interface show that CD45 is almost completely excluded from the membrane interface for both an anti-biotin IgG opsonized lipid (DPPE-biotin) and Fib1L antigen (Figure 5B). However, interfaces formed with Fib3L antigens were populated by freely diffusing CD45. Similarly high levels of CD45 were observed at the interface for both Fib5L and Fib7L antigens. Co-localization analysis revealed a size-dependent segregation threshold, with antigens Fib3L (estimated height 10.5 nm) and taller failing to exclude CD45 (Figure 5C).

We have previously shown that proteins >5 nm taller than a membrane interface are excluded from the reconstituted membrane interfaces due to steric interaction with the membranes (Schmid et al., 2016). To determine if our experimental results are consistent with exclusion of CD45, we computed the size of each Fibcon-family antigen-IgG-FcγR complex by adding the extended height of the antigen to the distance between the base of FcγR and the antigen-binding site of an IgG antibody (11.5 nm) (Lu et al., 2011), and compared this to the height of CD45RO, which is the sole isoform expressed in RAW 264.7 cells (PDB 5FMV, 22.5 nm) (Chang et al., 2016) (Figure 5D). The membrane interface formed by cells binding to opsonized lipid (estimated distance 11.5 nm) and opsonized Fib1L (estimated distance 15 nm) are both significantly shorter than CD45RO and are thus predicted to exclude CD45RO from the interface. However, the height of the interface formed by Fib3L (estimated distance 22 nm) is extremely close to the height of CD45RO and is therefore not expected to significantly exclude CD45RO. The Fib5L interface (estimated distance 29.0 nm) and Fib7L interface (estimated distance 36.0 nm) are both larger than CD45RO and are therefore not predicted to exclude CD45RO. These calculations are entirely consistent with the GPMV data and support a size-dependent mechanism of CD45 segregation that is governed by the antigen-mediated height of the membrane interface.

**Figure 5.**
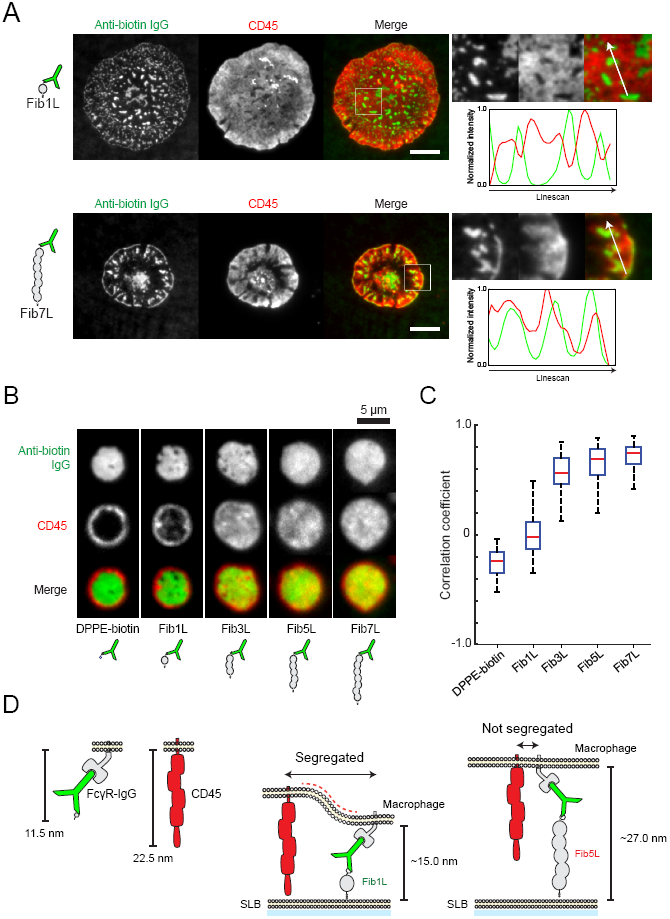
Phosphatase exclusion from antibody-FcγR clusters is antigen-height dependent. (A) Live-cell TIRF microscopy (100x) images at the macrophage-SLB contact interface for SLBs bound with Fib1L and Fib7L opsonized with anti-biotin IgG. For Fib1L SLBs (top), anti-biotin IgG (green, Alexa Fluor 488) is clustered at the cell-SLB interface, while CD45 (red, anti-CD45 Alexa Fluor 647) is segregated from IgG clusters. A line-scan through a region of the interface shows an anti-correlation between anti-biotin IgG and CD45 localization. For Fib7L SLBs (bottom), anti-biotin IgG is similarly clustered at the cell-SLB interface, however CD45 is not segregated from high-intensity IgG clusters. A line-scan through a region of the interface shows correlation between anti-biotin IgG and CD45 localization. Scale bar is 15 μm. (B) TIRF microscopy (100x) images at the GPMV-SLB contact interface for SLBs bound with DPPE-biotin, Fib1L, Fib3L, Fib5L, and Fib7L antigens. Anti-biotin IgG (green, Alexa Fluor 488) is enriched at the contact interface for each antigen. CD45 (red, anti-CD45 Alexa Fluor 647) is excluded in a size-dependent manner from DPPE-biotin and Fib1L, but not Fib3L, Fib5L, and Fib7L, GPMV-SLB contacts. Scale bar is 5 μm. (C) Quantification of the Pearson’s correlation coefficient between anti-biotin IgG (green) and anti-CD45 (red) channels for individual GPMV-SLB contacts. CD45 segregation, corresponding to a Pearson’s correlation coefficient of ∼0, is evident for DPPE-biotin and Fib1L antigens, but not for Fib3L, Fib5L, and Fib7L GPMV-SLB contacts. Box and whiskers denote inner quartile range and full range excluding outliers (> 1.5 quartile range). (D) Model of size-dependent segregation of CD45 at contact sites formed by FcγR-IgG binding. The FcγR-IgG complex spans ∼11.5 nm, while CD45RO is ∼22.5 nm tall. Since the membrane-membrane distance enforced by the FcγR-IgG-Fib1L complex spans ∼15.0 nm, CD45 is physically segregated from FcγR at the contact site in order to reduce membrane bending. In contrast, the FcγR-IgG-Fib5L complex spans ∼27.0 nm, which enforces a membrane-membrane gap that is large enough to accommodate CD45 at the contact site.

### Truncation of the CD45 ectodomain using CRISPR/Cas9 disrupts phagocytosis

If segregation of CD45 is necessary for FcγR phosphorylation and phagocytosis, then we expect that decreasing the size of the CD45 ectodomain would decrease phagocytic efficiency against short antigens. To test this without altering the endogenous balance of phosphatase and kinase activity and to avoid overexpression artifacts, we developed a strategy for truncating the ectodomain of endogenous CD45 using CRISPR/Cas9 (Figure 6A and Figure S4). Using two independent guide RNAs, we targeted the intronic region downstream of the first coding exon (containing the start codon and the signal peptide) and a separate intronic region directly upstream of an exon coding for the D3 FNIII domain. Excision followed by repair via non-homologous end joining resulted in a gene coding for CD45 protein with a truncated ectodomain, containing only the two final FNIII domains D3-D4. The ectodomain of this protein (CD45 D3-D4) is predicted to be 7.0 nm tall (based on analysis of full-length CD45, 5FN6), which is short enough to disrupt segregation of CD45 at a contact-interface formed through Fib1L.

We then compared the phagocytosis of a population of CD45 D3-D4 cells to wild-type CD45 cells. Truncation of the CD45 ectodomain significantly reduced phagocytosis of the short Fib1L antigens opsonized with anti-biotin IgG (Figure 6B-C). Microscopy images of cells collected during the process of phagocytosis show multiple beads bound to the cell periphery of the truncated CD45 D3-D4 cells, with dramatically reduced internalization relative to the full-length CD45RO cells, consistent with disrupted phagocytosis. Taken together, our results show that antigen size critically determines the success of antibody-dependent phagocytosis through size-dependent CD45 exclusion leading to FcγR phosphorylation. By differentially segregating kinase and phosphatase activity during interactions with opsonized targets, macrophages translate physical close contact into biochemical recognition and downstream activation.

**Figure 6.**
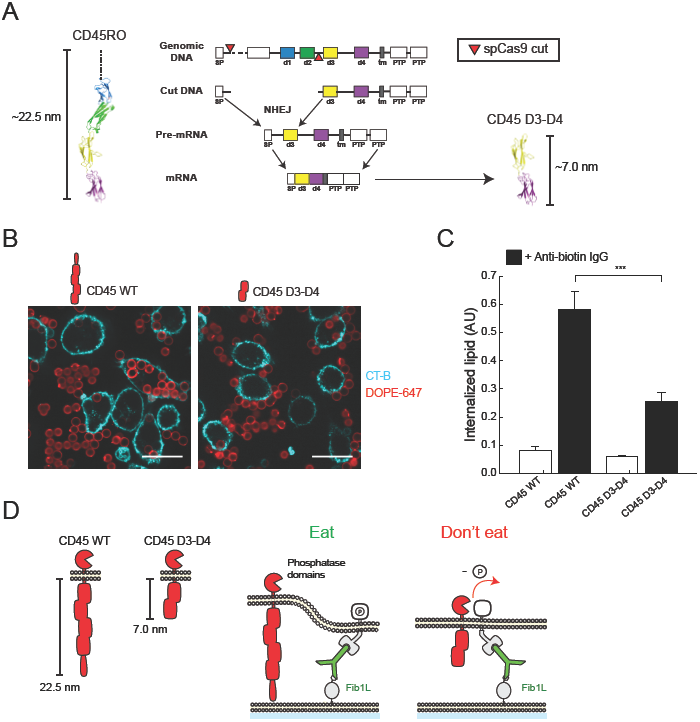
Truncation of the CD45 ectodomain using CRISPR/Cas9 disrupts phagocytosis. (A) Truncation of the CD45 ectodomain using CRISPR/Cas9 genome editing. Two independent guide RNAs that cut within intronic regions flanking the first coding exon and the exon coding for the d3 FNIII domain of CD45 were transduced into RAW 264.7 cells expressing spCas9. Dual cutting results in excision of the genomic region coding for the variable mucin domain and d1-d2 FNIII domains of CD45. Repair by non-homologous end-joining results in a gene coding for CD45 with a truncated ectodomain, containing only the D3-D4 FNIII domains, followed by the native transmembrane domain and tandem phosphatase domains of CD45. (B) Confocal fluorescence images (60x) of CD45 wild-type RAW 264.7 cells (left) and CD45 D3-D4 RAW 264.7 cells (right) incubated with Fib1L antigen and anti-biotin opsonized target particles. Scale bar is 20 μm. (C) Quantification of phagocytosis for CD45 wild-type and CD45 D3-D4 cells. Error bars are standard error over 3 independent wells. For each well, internalized lipid is an average quantification of N > 250 cells. P-values are paired Student’s T-test, where ***P < 0.001. (D) A model of the inhibition of phagocytosis by CD45 D3-D4. Truncation of the CD45 ectodomain reduces its height from ∼22.5 nm to ∼7.0 nm. An interface formed by FcγR-IgG-Fib1L binding spans ∼15.0 nm, which is sufficiently close to segregate wild-type CD45, reducing the local concentration of the phosphatase at sites of FcγR-IgG and triggering phosphorylation and activation of the macrophage. However, CD45 D3-D4 is ∼8 nm shorter than the interface, and thus is not segregated from the contact site. The failure to segregate CD45 D3-D4 upon FcγR-IgG binding leaves a higher local concentration of CD45 phosphatase at the contact site, suppressing phosphorylation and inhibiting phagocytosis.

## DISCUSSION

Antibodies have become vital therapeutic agents for battling cancer, autoimmune diseases, and neurodegenerative diseases. As a result, understanding how antibodies direct immune effector cell function, and how they might be targeted to promote desired behavior, has become increasingly important. Previous work has shown that biochemical properties of the antibody Fc domain, such as isotype and glycosylation, can produce divergent immune responses due to changes in binding specificity and affinity for activating and inhibitory Fc receptors (Jefferis, 2009; Nimmerjahn and Ravetch, 2005). However, less is known about how the physical properties of the target antigen, which anchors the antibody in a specific position and orientation relative to the cell surface, regulate the activation of Fc receptors and the response of immune effector cells. Given the physical diversity of potential target antigens, understanding mechanistically how the antigen-antibody-Fc-receptor complex triggers effector cell function is important for understanding fundamental principles of macrophage target recognition, interpreting different clinical responses to therapies, and designing more effective antibodies.

In this work, we demonstrate that antibody-dependent phagocytosis is controlled by the height of an antibody above the target-cell surface, which affects both Fc-receptor accumulation and phosphorylation. We engineered a family of synthetic antigens (Fibcon-repeat proteins) and used a family of tumor antigens (CEACAM) of different lengths to quantify changes in phagocytosis as a function of antigen size and to examine size-dependent phosphatase segregation that controls FcγR phosphorylation. We find that antigens that position an antibody <10 nm from cell-surface are optimal for stimulating phagocytosis. As a macrophage engages with a target surface, close contact stabilized by accumulation of receptors at the interface physically drives reorganization of signaling proteins on the macrophage cell surface. In particular, tall proteins are excluded by the small membrane-membrane gap while FcγR bound to antibody-opsonized antigens are enriched.

We demonstrate that the tyrosine-phosphatase CD45, which has a long ectodomain (∼22 nm), is excluded from sites of FcγR-antibody binding for antigens <10 nm tall, and that failure to segregate CD45 from interfaces formed by antigens >10 nm in height causes a reduction in FcγR phosphorylation. To confirm this finding, we show that reducing the size of the endogenous CD45 ectodomain with CRISPR/Cas9 disrupts phagocytosis against short antigens despite close membrane contact, proving that segregation of inhibitory phosphatases is required for phagocytosis. We believe this mechanism is analogous to the kinetic segregation model of T-cell activation (Shaw and Dustin, 1997; Wild et al., 1999), where elegant studies reconstituting the T-cell signaling network have provided evidence that the size-dependent segregation of the phosphatase CD45 from the TCR complex is necessary for T-cell activation (Carbone et al., 2017; Chang et al., 2016; Choudhuri et al., 2005; James and Vale, 2012). However, while TCR activation occurs through a receptor-ligand complex with defined binding geometry and characteristic height (15 nm) (Davis and van der Merwe, 2006), the height of the antigen-antibody-FcγR complex depends on both antigen height and antibody binding site, and thus naturally span a range of membrane-interface distances. How the variability in antibody height above the cell surface impacts immunosurveillance of tumor cells, which have been shown to upregulate large glycoproteins (Paszek et al., 2014), will be an interesting future area of study.

Our work demonstrates that a compact antigen-antibody-FcγR complex is sufficient to drive segregation of large inhibitory phosphatases and initiate macrophage phagocytosis, without the need for additional adhesion between the macrophage and target.

Interestingly, this differs from the recent observations of Freeman et al. (Freeman et al., 2016), who found that integrins and the cytoskeleton form a diffusion barrier that effectively segregates CD45 phosphatase to trigger phagocytosis. Interestingly, the model target surface used by Freeman et al. was a layer of immobilized antibody on a surface. In this situation, the density of FcγR that can accumulate within a given region is limited by the starting surface density of the antibody. However, if both the antibody and FcγR are capable of diffusion, as is the case for cell-surface tumor antigens such as CD20 and Her2 (though not for some pathogens), the local density of the FcγR can increase significantly at the membrane interface, with an upper limit set by steric constraints. In our minimal target system, where the antigen is free to move, we see such an increase. Our results suggest that enrichment of antibody-FcγR at contact sites on a fluid membrane is sufficient to activate phagocytosis, without requiring integrins to bridge low-density points of contact.

Our finding that size-dependent segregation is required for macrophage effector function offers an explanation for recent results with natural killer (NK) cells, which showed that antibody-dependent cellular cytotoxicity (ADCC) is most efficient for antibodies bound close the membrane (Cleary et al., 2017). Using NK cells, which activate ADCC via FcγR, Cleary et al demonstrated that the level of ADCC against target cells stimulated by the clinical monoclonal-antibody rituximab is high when it is bound to antigens that are < 8 nm tall but drops significantly when it is bound to an antigen that is ∼16 nm (CD37 tandem-repeat). Our results predict that ADCC should fail for a ∼16 nm antigen because the rituximab-antigen-FcγR complex forms a membrane interface that is tall enough to accommodate the phosphatase CD45, resulting in sustained dephosphorylation of the FcγR. However, the same study did not find a clear reduction in phagocytosis by bone-marrow derived macrophages (BMDM) against target-cells expressing the ∼16 nm tall antigen and opsonized with rituximab. One possible explanation for this is that the BMDMs interacting with the cellular targets used in that study engage a diverse set of pattern-recognition receptors (PRRs) in addition to FcγR (Gordon, 2002), all of which are capable of stimulating phagocytosis as well as altering the membrane interface gap size. Mechanistic details for different PRRs, including possible segregation height thresholds, could be isolated and studied using the reconstituted cell-surface like target particles presented here. In future work, we plan to explore whether and how combinations of PRRs engaged by a target work together or in opposition to determine immune effector cell function.

Use of reconstituted target particles in this study provides precise control over antigen physical properties, such as height and antigen density on the target membrane, making it possible to directly compare phagocytosis against different antigens. By combining these controlled targets with the display of fluid FcγR on the surface of GPMVs derived from macrophages, we found that for identical antibodies, increasing antigen height decreased the enrichment of antibody-FcγR complex, consistent with a decrease in affinity as predicted in physical models of receptor-ligand binding (Wu et al., 2011). Though this observation of reduced antibody affinity is not responsible for the decrease in FcγR phosphorylation with increasing antigen height (Figure 3D and E, and Figure S1E), reduced affinity could affect macrophage phagocytosis in several important ways. Beyond decreasing antibody-FcγR complex enrichment at the contact site, which reduces total ITAM available for phosphorylation, decreased affinity lowers the probability of initial adhesion and engagement with a target, weakens adhesion between membranes, and reduces the effectiveness of size-dependent phosphatase segregation. The effects of decreased affinity can be at least partially compensated for by increasing the surface density of antibody, which leads to increased phagocytosis for all antigen heights (Figure 2C).

While antibody-dependent phagocytosis of beads, cells, and pathogens clearly relies on dynamic assembly of the actin cytoskeleton for engulfment, we found that size-dependent phosphatase segregation and FcγR phosphorylation did not require an intact actin cytoskeleton. This suggests that early signaling events depend largely on local organization of transmembrane and membrane-bound components, while dynamic actin network assembly controls late phase spatial reorganization of cell-surface components (Jaumouillé and Grinstein, 2011) and ultimately particle internalization. We suggest that size-dependent segregation of proteins at membrane interfaces is a central organizing principle that is always influencing FcγR phosphorylation, but it is superposed with other influences on lateral membrane organization that could affect clustering and phosphorylation state, including the actin cytoskeleton and lipid microdomains (Beekman et al., 2008; Lopes et al., 2017). How these different physical mechanisms are coordinated, both in time and in space, is a subject of interest for future studies.

There is increasing evidence that the location of antibody binding epitopes is important for the activity of antibody-dependent effector cell activity (Cleary et al., 2017). Remarkably consistent with our results, the majority of clinical antibodies that rely on direct effector-cell activity bind to relatively small cell-surface receptors (i.e. CD20 2.1 nm, EGFR 6.9 nm, HER2 3.8 nm, CTLA4 2 nm, PD-1 5.2nm, PD-L1 7.1 nm) (Gül and van Egmond, 2015) (Supplementary Table 1). It has previously been shown that antibodies binding close to the membrane display an enhancement in complement-dependent-cytotoxicity (CDC) cell killing, which may lead to increased activity of antibodies with surface-proximal epitopes (van Meerten et al., 2010). Our results suggest that similar enhancement of ADCP and ADCC for surface-proximal epitopes could occur through increased segregation of inhibitory phosphatases from regions of FcγR-antibody accumulation.

The size-dependent signaling reported here may have general implications for receptors with similar signaling networks in multiple immune-cell types. In addition to macrophages, T-cells and NK cells targeted to tumors through bi-specific antibodies and chimeric antigen receptors rely on selection of tumor antigens with appropriate physical properties to promote activation of signaling molecules. The results of our study suggest that this could be accomplished by tuning the membrane interface gap to physically exclude inhibitory molecules in order to trigger immune cell effector function.

## Acknowledgements

The authors would like to thank K. Heydari and M. West for help with flow cytometry and Fletcher Lab members for helpful feedback and technical consultation. This work was supported by the Immunotherapeutics and Vaccine Research Initiative (IVRI) as well as the NIH R01 GM114671 (DAF). M.H.B. was funded by an NSF and a Siebel Fellowship. S.S. was funded by an LSRF fellowship. D.A.F. is a Chan Zuckerberg Biohub investigator.

## EXPERIMENTAL PROCEDURES

### Chemical reagents

HEPES (4-(2-hydroxyethyl)-1-piperazineethanesulfonic acid), MOPS (3-(N-morpholino)propanesulfonic acid), TCEP (tris(2-carboxyethyl)phosphine), KCl, NaCl, glucose and sucrose were purchased from Fisher Scientific. Imidazole was purchased from Sigma Aldrich. DOPE-647 (1,2-Dioleoyl-sn-glycero-3-phosphoethanolamine labeled with Atto 647) was purchased from ATTO-TEC. POPC (1-palmitoyl-2-oleoyl-snglycero-3-phosphocholine), DGS-Ni-NTA (1,2-dioleoyl-sn-glycero-3-[(N-(5-amino-1-carboxypentyl)iminodiacetic acid)succinyl], with nickel salt) and DPPE-biotin (1,2-dipalmitoyl-sn-glycero-3-phosphoethanolamine-N-(biotinyl)) were purchased from Avanti Polar Lipids (Alabaster). All chemical reagents were used without further purification. Anti-biotin IgG (anti-biotin monoclonal mouse IgG1 antibody BK-1/39 labeled with Alexa Fluor 488) was purchased from ThermoFisher. Anti-CEA IgG (anti-pan CEACAM mouse monoclonal IgG1 antibody) was purchased from Santa Cruz Biotechnology. CMFDA (CellTracker Green), Hoechst 33342, and CT-B-555 (Cholera toxin subunit B with Alexa Fluor 555 conjugate) dyes were purchased from ThermoFisher. Lat-A (Latrunculin A) was purchased from Abcam.

### Preparation of minimal target cell (target particles)

Target particles were generated by coating glass beads with a fluid supported lipid bilayer, to which antigens with specific antibody binding sites were attached. The individual steps are described in detail below.

#### Preparation of small unilamellar vesicles (SUVs)

SUVs were prepared by rehydrating a lipid film composed primarily of POPC, doped with up to 2% of DPPE-biotin or DGS-NI-NTA and 0.8% DOPE-647 in pure deionized H20. The rehydrated solution was vortexed briefly, sonicated at low-power (20% of max) using a tip-sonicator, and finally filtered through a 200 nm PTFE filter (Millipore). Solutions of SUVs were stored on ice and used within 48 hours to avoid phospholipid oxidization.

#### Preparation of supported-lipid bilayer (SLB) coated glass beads

40 μL of 3.78 μm glass bead (Bangs labs) slurry (10% solids) were cleaned using a 3:2 mixture of H2SO4:H2O2 (Piranha), and clean beads were spun down at 1000 G and washed 3 times before being resuspended in 400 μL of pure water. Clean beads were stored in water at room temperature and used within 48 hours. To assemble supported-lipid bilayers, 20 μL of SUV solution was diluted in 80 μL of MOPS buffer (25 mM MOPS pH 7.4, 125 mM NaCl,), and 10 μL of clean bead slurry were added and mixed gently by pipetting. The bead/SUV mixture was incubated for 15 minutes at room temperature while rotating continuously to reduce bead sedimentation. Beads were spun down gently at 50 G for 1 minute, and SUV solution was carefully removed and replaced with PBS. The fluidity of the lipid-bilayer was assessed by imaging beads deposited on a glass coverslip with a spinning-disk confocal microscope (Nikon) at 60x magnification and high laser power, where diffusion of single-molecules of labeled lipid was visible after photo-bleaching a small region-of-interest.

#### Reconstitution of antibody-opsonized target particles

SUV mixtures with up to 2% DGS-Ni-NTA were used to prepare SLB-coated beads. To prepare target particles, beads were incubated with 50 nM of recombinant protein antigen containing a C-terminal 10-His tag for 15 minutes. The protein binds fluidly to the surface via the Nickel-His interaction, and the interaction of one-protein with up to ten DGS-Ni-NTA lipids lead to nearly irreversible attachment (Chikh et al, 2002). To prepare opsonized target particles, anti-biotin IgG was added at 1-5 ng/mL and incubated along with the protein, such that the anti-biotin IgG bound fluidly to the surface via interaction with the His-tagged protein.

### Preparation of size-variant protein antigens

Two families of proteins were prepared and bound to the supported lipid bilayer of the target particles to present an antibody at a known distance from the membrane.

#### Design of size-variant Fibcon repeat protein antigens (Fib1L-Fib7L)

The FNIII domain occurs with high frequency in cell-surface proteins, where it is often linked together in an N-C topology to create proteins with extended height (Doolittle et al, 1995). We designed a family of synthetic proteins that similarly rely on the FNIII domain to generate height. The Fibcon domain is a high-stability FNIII domain designed through multiple-sequence alignment (Jacobs, PEDS). The DNA sequence coding for the Fibcon protein was ordered as a synthesized gene fragment (Integrated DNA Technologies). Repeats of the Fibcon sequence were cloned into a pET28b vector (EMD Millipore) for expression in E. coli cells with no linker region via Gibson assembly (Gibson et al, 2009). The Fibcon repeat sequence was flanked by an N-terminal YBBR peptide (Yin et al, 2005) and a C-terminal His-10 followed by a KCK sequence for chemical labeling, and terminated with an additional His-6 sequence.

#### Fibcon family protein expression and purification

All proteins were expressed in Rosetta DE3 competent cells (EMD Millipore). Cells were grown at 37 °C to OD = 0.8, induced with 0.3 mM IPTG overnight at 18 °C. Cells were harvested and resuspended in 25 mM Hepes pH 7.4, 150 mM NaCl, 0.5 mM TCEP and 10 mM imidazole, and lysed by freeze thawing and sonication. The lysate was centrifuged for 45 min at 20,000g, affinity purified over a His-Trap HP column (GE Healthcare) through imidazole gradient elution on an AKTA Pure (GE Healthcare) system. Peak fractions were concentrated and gel-filtered via a Superdex 200 column into 25 mM HEPES pH 7.4, 150 mM NaCl, 0.5 mM TCEP. Proteins were concentrated, and purity was assayed on an SDS page gel. To compare Fib1L, Fib3L, Fib5L, and Fib7L proteins, the protein molecular weight was verified by an elution shift during gel-chromatography from a Superdex 200 column (Figure S1A).

#### Height measurement of Fibcon-family antigens

Fibcon proteins were fluorescently labeled at the N-terminus and conjugated to glass beads coated in a fluorescent lipid bilayer. The bead was immobilized on glass surface to minimize vibration during imaging. Confocal images of the bead’s equatorial plane were repeatedly captured while fluorescence of protein and lipid were being illuminated alternatively. Radial distance between the protein and lipid fluorescence were estimated in pixels based on multiple pairs of acquired images and converted to nm using known calibration. The radial offset represents the mean height of the protein. The measurement was repeated in 8 to 10 different beads and the mean and standard deviation were reported for each fibcon proteins.

#### Site-specific biotinylation of Fibcon family proteins with SFP synthase

Fibcon proteins were biotinylated at the N-terminus using an SFP synthase catalyzed reaction, which conjugates biotin to the YBBR tag. 100 μM recombinant Fibcon protein, 120 μM biotin CoA, 10 μM SFP synthase (purified according to published protocols (Yin et al, 2006)), and 40 mM MgCl were mixed in a 100 μL reaction volume and rotated for 3 hours at room temperature. The labeled protein product was purified on a Superdex 75 10/300 gel filtration column (GE Healthcare) into 25 mM Hepes pH 7.4, 150 mM NaCl, 0.5 mM TCEP. Biotinylation of the product was confirmed by attaching the protein to a supported lipid bilayer and imaging the binding of anti-biotin IgG.

#### Design and characterization of size-variant CEACAM5 protein antigens

Human CEACAM5 (Uniprot P06731) was chosen as a model antigen due to its relevance in cancers and relatively tall height of seven Ig-like domains (estimated ∼28 nm). Full length CEACAM5 (CEA-FL, AA 34-677) DNA was cloned into pHR lentiviral expression vector (Clonetech). In parallel, a shortened version of CEACAM5 (CEA-N, AA 34-144) containing only the N-terminal domain linked to the native GPI anchor was cloned into the same vector. Anti-CEA IgG was expected to bind to the N terminal domain of all CEACAMs, as this is the only domain shared by the family. To test this, CEA-FL and CEA-N were transiently transfected in HEK cells using TransIT-293T transfection reagent (Mirus Bio) along with a construct containing only the two membrane proximal domains of CEACAM5 as a control. These cells were fixed and binding of Anti-CEA IgG was confirmed in the case of both CEA-FL and CEA-N, but not in the control lacking the N-terminal domain (Figure S2).

#### Preparation of CEA-N and CEA-FL protein antigens

HEK293T cells were grown to 70% confluency in a T175 flask and transfected with CEA-N or CEA-FL DNA in a pCAGGS expression vector using TransIT-293T transfection reagent (Mirus Bio). After 48 hours, the supernatant was collected and Halt™ protease and phosphatase inhibitor was added (ThermoFisher). The proteins were affinity purified over a His Trap Excel column (GE Healthcare) and eluted with a high imidazole buffer containing 25mM HEPES, 150 mM NaCl and 500 mM imidazole at pH 7.4. The proteins were gel-filtered using a Superdex 200 column (GE Healthcare) and the buffer was exchanged to remove imidazole. The proteins were concentrated and the purity was confirmed using SDS-PAGE (Figure S2).

### Quantification of phagocytosis

Phagocytosis of target particles by macrophage-like RAW 264.7 cells was quantified using microscopy and flow cytometry.

#### Microscopy assay of phagocytosis

96-well flat-bottom tissue-culture plates (Corning) were seeded with 35,000 cells in 200 μL of RPMI 1640 medium. Cells were incubated at 37 °C for at least 2 hours to allow attachment to the plastic surface. To start the assay, 100 μL of target particle suspension containing ∼500,000 beads was added to each well, and the plate was returned to 37 °C for exactly 20 minutes. After 20 minutes, wells were washed once with PBS to remove non-internalized and partially bound beads, and then overlaid with a PBS containing 1 μM CMDFA and 10 μM Hoechst 33342 to fluorescently stain the cell-cytoplasm and cell-nuclei. Individual wells were imaged after 10 minutes with the staining solution on a spinning-disk confocal microscope (Nikon) at 20x. For each well, a grid pattern of 4 fields-of-view was recorded. Images were segmented using a routine written with CellProfiler (Broad Institute) to isolate single-cells, and the bead fluorescence intensity in the lipid (DOPE-647) channel was integrated on a single-cell basis to generate the quantification of internalized lipid.

#### Flow-cytometry assay of phagocytosis

96-well plates were prepared for phagocytosis as described above. To start the assay, 100 μL of bead-protein-antibody solution was added to each well, and the plate was returned to 37 °C for 20 minutes. After 20 minutes, wells were washed once with PBS to remove non-internalized and partially bound beads, and then overlaid with PBS containing 20 mM EDTA and 10 μM Hoechst 33342. Cells were gently de-adhered by pipetting up and down, then left suspended within the 96-well plate for flow-cytometry. Flow cytometry was performed on the Attune NxT equipped with an autosampler and analyzed with the provided software (Thermo Fisher). Single-cells were gated using the Hoechst channel in addition to forward and side-scatter. The 647 channel recording the fluorescence of DOPE-647 was used to quantify internalized lipid per cell (Figure S1D).

### Probes for imaging of FcR phosphorylation

The phosphorylation state of the FcR ITAM was detected using both immunofluorescence and a live-cell sensor based on Syk kinase.

#### Immunofluorescence of FcR phosphorylation

For imaging interfaces between cells and supported lipid bilayer coated beads, cells were seeded into 8-well imaging chambers with a cover-glass bottom (Cellvis) and beads were added to the wells once the cells had fully adhered to the cover-glass. After a 15 minute incubation at 37 °C, the cells were fixed for 10 minutes with 4% paraformaldehyde in PBS. Cells were permeabilized with 0.1% saponin (Alfa Aesar) and blocked with 3% (w/v) BSA in PBS along with 0.5 μg/mL Fc Block™ (BD Biosciences). Saponin (0.1%) was included in all subsequent probing and washing steps. Phospho-Tyrosine antibody (P-Tyr-1000 MultiMab™, Cell Signaling Technology) was added to cells at a dilution of 1:500 and incubated at room temperature for 1 hour. The cells were washed and secondary antibody (Alexa Fluor 488 AffiniPure Donkey Anti-Rabbit IgG, Jackson ImmunoResearch) was added at a dilution of 1:1000 and incubated for 1 hour at room temperature. The cells were given a final wash in PBS before imaging.

#### A live-cell sensor of FcR phosphorylation based on Syk kinase tandem-SH2 domains

The tyrosine-protein kinase Syk is recruited to the phosphorylated ITAM of Fc-receptors via an interaction with its tandem-SH2 domains (Chu et al, 1998). A sensor that localizes specifically to phosphorylated ITAMs was designed by placing a fluorescent protein C-terminal from the isolated Syk SH2 domains. The sensor construct consists of a C-terminal mCherry fluorescent protein, followed by a linker region (GGGSGGGG), followed by amino acids 2-261 of the tyrosine-protein kinase Syk from Mus musculus (NP_035648), a region which covers the tandem SH2 domains of Syk (Figure S3B). The sensor was cloned into the pHR lentiviral expression vector (Clonetech) under control of the low-expression UBC promotor (reference).

#### Preparation of lentivirus with the live-cell sensor of FcR phosphorylation

HEK293T cells were grown in a 6-well plate to 80% confluency, and 160 ng VSV-G, 1.3 μg CMV 8.91, and 1.5 μg target vector were transfected into HEK293T cells using TransIT-293T transfection reagent (Mirus Bio). Viral supernatants were collected 60 hours after transfection and spun at 4000 G to remove HEK293T cells. Viral supernatant was stored at 4 °C for no longer than 48 hours prior to infection. For lentiviral infection, 500 μL of viral supernatant was added to 5e5 RAW 264.7 macrophages along with 4 μg/mL polybrene, and cells were spun at 400G for 25 minutes at 37 °C and then resuspended and plated in a 6-well plate. Viral media was replaced fresh growth media 24 h after infection. Cells were sorted via fluorescence-activated cell sorting on an Influx Cell Sorter (Beckton-Dickinson), and a population of cells expressing the mCh-Syk-SH2 sensor was expanded and frozen for later use.

### Live cell imaging and quantification of FcR phosphorylation

FcR ITAM phosphorylation of macrophages engaged with antigens on a supported lipid bilayer was imaged in TIRF microscopy.

#### Preparation of SLB on coverslips

SLBs were formed by fusion of SUVs (see ‘Preparation of small unilamellar vesicles (SUVs)’) to RCA-cleaned glass coverslips. 40 μL of SUV solution was diluted in 60 μL of MOPS buffer (25 mM MOPS, 125 mM NaCL, pH 7.4) in a PDMS chamber sealed over an RCA cleaned coverslip. The SUV mixture was incubated for 15 minutes at room temperature. Next, the excess SUVs were thoroughly removed by washing 5x with 60 μL of PBS without drying the coverslip. The fluidity of the resulting lipid-bilayer was assessed by imaging with a spinning-disk confocal microscope (Nikon) at 60x magnification and high laser power, where diffusion of single-molecules of labeled lipid was visible after photo-bleaching a small region-of-interest.

#### Live-cell Total Internal Reflection (TIRF) microscopy of FcR phosphorylation

A solution consisting of 50 nM antigen protein and 5 ng/mL of IgG antibody was added to the hydrated SLB and incubated for 15 minutes at 37 °C. 1e4 RAW 264.7 were added dropwise to the imaging chamber and allowed to settle towards the SLB over 5 minutes. Where stated, cells were preincubated with CT-B-555 to stain the cell membrane. TIRF imaging was performed on a Ti Eclipse microscope (NIKON) using a 60x TIRF 1.49 NA objective and an iXon Ultra EMCCD (Andor). All imaging experiments were performed within an incubator stage insert in a 5% CO2 environment at 37 °C (Oko labs).

#### Image processing of FcR phosphorylation

Images were processed with custom code written in Python (Python.org) and MATLAB (MathWorks). For quantifying mCh-Syk-SH2 localization at the plasma membrane, single-cells were segmented using the CT-B-555 channel, and the intensity of the mCh-Syk-SH2 channel was integrated within this region to quantify sensor-recruitment. To quantify mCh-Syk-SH2 localization to individual clusters of antibody-FcR, Otsu thresholding was performed to isolate high-intensity anti-biotin IgG clusters from the background level of anti-biotin IgG bound to the SLB, and the intensity of the mCh-Syk-SH2 channel within these clusters was averaged to quantify sensor recruitment for each cell.

#### Live-cell antibody-based TIRF imaging of CD45 localization

To image CD45 localization, cells were incubated with 0.5 μg/ml anti-mouse CD45 antibody (clone 30-F11) directly conjugated to Alexa Fluor 647 (Biolegend) for 10 minutes. 50 μL of cells (1e4 cells) were diluted directly into 100 μL imaging chambers containing hydrated, protein and antibody bound SLBs (described above). After allowing cells to settle to the SLB over 5 minutes, two-color TIRF images of CD45 (647) and anti-biotin IgG (488) localization were collected on newly surface-engaged cells over a period of 15 minutes.

### Preparation and use of giant plasma-membrane vesicles

The plasma membrane of RAW 264.7 cells was isolated and used to quantify affinity of antibody-bound antigens to FcR and segregation of CD45.

#### Giant plasma membrane vesicle (GPMV) formation

GPMVs were made following the protocol outlined by Sezgin et al. (Sezgin et al, 2012). In brief, cells were seeded in a 6-well plate and allowed to adhere. They were then washed with buffer (10 mM Hepes pH 7.4, 150 mM NaCl, 2 mM CaCl2) before addition of vesiculation agent (25 mM PFA and 2 mM DTT in the same buffer). GPMVs formed for one hour at 37 °C and were collected by removing the supernatant from the cells.

#### TIRF imaging of GPMV-SLB interfaces

GPMVs were added directly to 100 μL imaging chambers containing hydrated, anti-biotin IgG opsonized SLBs (described above). After settling for 15 minutes, FcR-engaged GPMVs were identified by enrichment of the anti-biotin IgG signal and TIRF images of the GPMV-SLB interfaces were captured. To image CD45 localization, Alexa Fluor 647 labeled CD45 antibody (clone 30-F11) (Biolegend) was added to the well to a concentration of 0.5 μg/mL. CT-B-555 was used as a general membrane stain for GPMVs and was added to the well at a concentration of 0.5 μg/mL. The GPMVs were imaged after an incubation period of 10 minutes.

#### Enrichment analysis

For quantifying antibody-FcR enrichment at the GPMV-SLB contact, single GPMV footprints were segmented using the CT-B-555 channel, and the intensity of the anti-biotin IgG channel was averaged within this region to quantify antibody intensity within footprint region (‘in’). The average intensity within the background region (‘out’) was measured, and an ‘enrichment ratio’ for each footprint was calculated as the ratio of intensity in region ‘in’ over region ‘out’ (Figure 4b).

#### Correlation analysis

For quantifying correlation between CD45 and anti-biotin IgG, the CT-B-555 channel was used to segment regions of GPMV-SLB contact. The Pearson’s correlation coefficient was calculated between the CD45 and anti-biotin IgG channels within these regions to quantify colocalization.

### CRISPR editing to generate CD45 truncation

The gene coding for the CD45 protein from Mus musculus (geneID 19264) was truncated by a CRISPR/Cas9 exon excision strategy (Nelson et al, 2015) using guide RNAs targeting the intronic regions preceding the second coding exon (Exon 3) and directly following the exon coding for the D3 FNIII domain (Exon 8), resulting in a gene coding for CD45 protein with a truncated ectodomain containing only the two final FNIII domains D3-D4 (CD45 D3-D4) (Figure 6A).

#### sgRNA design and expression vector cloning

Target sequences for wild-type spCas9 nuclease were selected using the Deskgen CRISPR/Cas9 design tool (www.deskgen.com). For each genomic region to be cut, two target sequences were selected for high activity scores via the algorithm in Doench 2016 and low chance of off-target effects as calculated by the algorithm in Hsu et al 2013. For the region preceding Exon 3 (Chr1 bp 138126429 – 138126489), two target sequences were selected: Start-1 CTAATGGATGACCTAAGATG TGG, Start-2 AGAGCAATTCCTGTAACGGG AGG. For the region following Exon 8 (Chr1 bp 138110583 – 138110643), two target sequences were selected: D3-1 AAACCTTATTAAATAGAAAG GGG, D3-2 TGGTGTTATAAAAAGAAGGG AGG. Oligos coding for single guide RNAs (lacking the PAM sequence) were purchased from Integrated DNA Technologies (IDT) and cloned into the lentiGuide-Puro plasmid using the BsmB1 restriction enzyme as previously reported (Sanjana et al, 2014).

#### Transduction of spCas9 into RAW 264.7 cells

Lentivirus was generated from the lentiCas9-Blast plasmid in HEK293T cells and transduced into RAW 264.7 cells as described above. After 2 days, 5 ug/mL Blasticidin was added to the media to select for Cas9 expressing RAW264.7 cells. Cells were cultured in 2 ug/mL Blasticidin for two weeks before freezing in 90% FBS + 10% DMSO for later use.

#### Transduction of sgRNA plasmids into RAW 264.7 cells to trigger genomic excision

Lentivirus was generated from lentiGuide-Puro plasmids encoding the Start-1, Start-2, D3-1, and D3-2 guide RNAs as described above. To excise the genomic region coding for the mucin-like and D1-D2 FNIII domains of CD45, two guide RNAs were transduced into Cas9-expressing RAW 264.7 cells simultaneously, one targeting the Exon 2 region (Start) and one targeting the Exon 8 region (D3). Four cell-lines were transduced using pairs of lentivirus (Start-1+D3-1, Start-2+D3-2, Start-2+D3-1, Start-2+D3-2). After 2 days, media containing 2 ug/mL Puromycin and 5 ug/mL Blasticidin was added to the media to select for cells expressing both Cas9 and at least one sgRNA cassette. Cells were cultured for 2 weeks for recovery. In a subset of the cells where two guide RNAs were transduced, simultaneous cutting by spCas9 at two genomic sites will lead to the removal of a large genomic region (∼16,000 bp), which in a further subset of cells will be repaired by non-homologous end-joining.

#### Selection of CD45 D3-D4 cells

To identify cells expressing CD45 D3-D4 we used the monoclonal antibody anti-CD45 (clone 30-F11), which binds to an epitope in the pan-CD45 cysteine-rich domain D1 (Symons et al, 1999). Cells were collected two weeks after sgRNA transduction and antibiotic selection, labeled with anti-CD45 (clone 30-F11) Alexa Fluor 647, and cells were sorted by anti-CD45 labeling on a Bioscience Influx Sorter (BD) (Figure S4). Two populations of cells were recovered, CD45 30-F11 positive and CD45 30-F11 negative. After growing for one week, CD45 30-F11 negative cells were again labeled and sorted to remove any remaining positive cells from the population. At this stage, the resulting CD45 30-F11 negative population may contain cells expressing the truncated CD45 D3-D4. Two populations of cells, CD45 30-F11 positive CD45-WT and CD45 30-F11 Negative (CD45 D3-D4), were frozen in 90% FBS + 10% DMSO for later use.

#### RT-PCR to detect CD45 D3-D4 mRNA

To characterize and confirm the CD45 truncation in macrophages, the RNA was extracted from cells and then reverse transcribed to DNA and amplified. RNA extraction from whole cell lysate was performed with the RNeasy Mini kit (Qiagen). Once extracted, the RNA was reverse transcribed and amplified using the OneTaq RT-PCR kit (New England BioLabs). During this step, CD45 RNA was specifically targeted using primers complementary to it beginning near the 5’ end (gctgatctccagatatgaccatggg) and ending near the transmembrane encoding region (gacatcaatagccttgcttgttgttttgtat). This process of RNA extraction and RT-PCR was performed on cells edited to contain only the D3-D4 domains of CD45 (D3-D4) as well as wild-type cells (WT). The amplified DNA from all cells was assayed on an agarose gel to determine length and the CD45 D3-D4 sample was clearly smaller than the WT sample – indicating that this population of cells had a truncation in CD45 mRNA length (Figure S4B). The individual bands from each population were excised and the DNA was sequenced to further confirm the CD45 truncation in cells (Figure S4C).

#### Western Blot Analysis to confirm CD45 protein truncation

Cells were grown to 80% confluency in a T175 tissue culture flask. The cells were mechanically lysed using a dounce homogenizer and fractionated to isolate the plasma membranes. The membrane fraction was ultimately suspended in HEPES buffer with detergent (25 mM HEPES, 150 mM NaCL, 0.5% NP-40). Protein concentrations were measured via BCA assay to ensure equal amounts of sample were separated with SDSPAGE. After transferring the protein to a nitrocellulose membrane, the membrane was probed with anti-CD45 antibody (PAB030Mu01, Cloud-Clone Corp.) at a dilution of 1:400. This antibody was chosen because it was raised against the D3-D4 domains of CD45, which are present in both full length CD45 and the truncated variant. After washing, the membrane was probed with anti-rabbit HRP secondary antibody (ab6721, Abcam) at a 1:5000 dilution.

---

**Supplementary Video 1**

**Confocal fluorescence microscopy video of a RAW 264.7 cell spreading on an IgG opsonized supported-lipid bilayer bound with Fib1L antigen.** Fluorescence signal is Alexa Fluor-488 labeled anti-biotin IgG attached to the bilayer through Fib1L antigen, and enrichment over background corresponds to binding to FcgR at the apposed cell footprint and subsequent immobilization. Time course for the video is 2 minutes 25 seconds, and the scale bar is 10 μm.

**Supplementary Video 2**

**TIRF microscopy video of a RAW 264.7 cell expressing pITAM sensor spreading on an IgG opsonized supported-lipid bilayer bound with Fib1L antigen.** Alexa Fluor-488 labeled anti-biotin IgG attached to the bilayer through Fib1L antigen (left), corresponding to bound antibody-Fcgr complexes, and pITAM sensor within the cell (center) are co-localized at the contact interface. Alexa Fluor-488 (green) and mCherrypITAM (red) channels are merged to highlight co-localization (right). Time course for the video is 2 minutes 25 seconds, and the scale bar is 10 μm.

